# Past, present, and future of the Living Planet Index

**DOI:** 10.1101/2022.06.20.496803

**Authors:** Sophie E H Ledger, Louise McRae, Jonathan Loh, Rosamunde Almond, Monika Böhm, Christopher F Clements, Jessica Currie, Stefanie Deinet, Thomas Galewski, Monique Grooten, Martin Jenkins, Valentina Marconi, Brett Painter, Kate Scott-Gatty, Lucy Young, Michael Hoffmann, Robin Freeman

**Author notes:** Joint first and corresponding authors:.

## Abstract

As we enter the next phase of international policy commitments to halt biodiversity loss (e.g. Post-2020 Biodiversity Framework), biodiversity indicators will play an important role forming the robust basis upon which targeted, and time sensitive conservation actions are developed. Population trend indicators are perhaps the most powerful tool in biodiversity monitoring due to their responsiveness to changes over short timescales and their ability to aggregate species trends from global down to at a sub-national or even local scale. We consider how the project behind the foremost population level indicator - the Living Planet Index - has evolved over the last 25 years, its value to the field of biodiversity monitoring, and how its components have portrayed a compelling account of the changing status of global biodiversity through its application at policy, research and practice levels. We explore ways the project can develop to enhance our understanding of the state of biodiversity and share lessons learned to inform indicator development and mobilise action.

**Box 1. The Living Planet Index Project**
The Living Planet Index project (the index, methodology, and database) and its secondary outputs (methods papers and R code, database and website, global index, and subset indices) have had wide-ranging applications within the fields of biodiversity monitoring and research, as well as across policy, education, and outreach.
The Living Planet Index (LPI) is a biodiversity indicator which tracks trends in the relative abundance of wild vertebrate populations (where population is defined as to a single species in a defined location rather than the biological definition). Relative abundance captures how populations are changing over time on average in comparison to a reference point, or “baseline” (the LPI uses 1970). It is often described as analogous to a stock market index for species. The index is comprised of thousands of population time-series for vertebrate species from locations around the world; the trends from these populations are averaged to produce terrestrial, freshwater, and marine indices, which are further aggregated to a global LPI. The latest global LPI shows a decline of 68% between 1970 and 2016 globally ^1^. This is an average trend based on time-series data from 20,811 populations of 4,392 species of mammals, birds, reptiles, amphibians, and fish.
The LPI database (LPD) can include population data for any species for which time-series population data could be found, regardless of threat status, or whether they show increasing or declining trends. These population time-series are sourced from scientific papers, online databases, government, and expert led published reports. They can be searched and downloaded from the project website (www.livingplanetindex.org). More technical information is available on the LPI stats website (http://stats.livingplanetindex.org/).

## Introduction

The Living Planet Index (LPI) (Box 1) was first proposed as a means of evaluating environmental change, particularly by tracking trends in global biodiversity, a quarter of a century ago ^2^. At that time, although there was mounting evidence of anthropogenic impacts on nature ^3^, there were very few indicators of the state of biodiversity or ecosystems at a global, or even regional scale. The initial version of the LPI, based on trends in vertebrate populations and forest cover, indicated that biodiversity was in decline globally ^2^. A successful response to what is now widely recognised as a global biodiversity crisis ^4-7^ will involve transformative changes in the way humans use the planet’s resources,^8-10^ widespread intergovernmental action ^11^ and ambitious targets ^9,10^ (intergovernmental agreements such as the Convention on Biological Diversity (CBD) ^12^ and the United Nations Sustainable Development Goals (SDGs) ^13^). To this end we need meaningful and reliable biodiversity indicators, generated from high quality and large-scale data to track progress towards targets down to the national level ^10,14^. As such, the development of biodiversity indicators has become an increasing focus in conservation science ^15-17^, particularly to ensure they are fit for purpose as tools for management and policy, as well as to improve the representation of the underlying data beyond well-studied taxa and regions.

Within this review we chart the history, progression, and applications of the LPI project (Box 1). We review the LPI as a tool for public engagement and outreach, policy, and to drive further research and, analyse citation data to explore other applications of the LPI. We discuss challenges faced in maintaining a large biodiversity dataset and in current uses of the LPI. Finally, we look to the future and propose how the LPI project could evolve by enabling global collaboration to strengthen the indicator, harnessing new technologies for collecting population data, and developing new analysis to better understand the relationships between drivers and wildlife population trends.

### The origins and development of the Living Planet Index

The Living Planet Index was conceived in 1997 by the World Wildlife Fund for Nature (WWF International). The primary aim was to *“develop a measure of the changing state of the world’s biodiversity over time*” ^18^ using aggregate time-series population trends for a large sample of species from across the world. As very little data were available on plants, fungi or invertebrate species, the pragmatic approach was taken to restrict the initial LPI taxonomically to vertebrates. There was also geographic unevenness in the distribution of the available data: long-term monitoring studies dating back decades were located mainly in Europe and North America. To address the biases in data coverage, a benchmark of 1970 was set, and the data were divided up into three broad biomes – terrestrial, freshwater and marine – and then further into regional groupings. The source data and LPI outputs were at first collaboratively managed by WWF and the World Conservation Monitoring Centre (now UN Environment Programme WCMC) for use within WWF’s flagship publication, the Living Planet Report (LPR). First published in 1998, the LPR used the initial iteration of the LPI as a communications tool to convey biodiversity trends into a singular message on the health of the planet for a broad audience, alongside measures of humanity’s impact on the planet ^2^. Calculated as -32% between 1970 and 1995 (Loh, et al. ^2^), the downward trend of the LPI was already apparent.

In the early 2000’s, as the LPI dataset and methods were developed further ^18^, their potential for use in advocacy, research, and as an indicator for monitoring biodiversity were recognised more widely. In 2002, the Parties to the CBD committed to achieve a significant reduction of the rate of biodiversity loss at the global, regional and national level by 2010 and required a framework of biodiversity indicators to monitor their progress ^19^. The first national LPI, the ‘Living Uganda Index’, was published with the National Biodiversity Data Bank recording scheme at Makerere University, Uganda in 2004 ^20,21^ and was presented as a case study for country-level applications of species population indices at CBD COP 7 ^22^. A Discussion Meeting held at the Royal Society in 2004 brought together leading academic and NGO researchers working on biodiversity indicators, and the resulting papers, including one on the LPI, were published in a special issue of Philosophical Transactions B ^23^. This meeting laid much of the groundwork for subsequent indicator development in the context of the CBD and other international biodiversity monitoring processes ^24^. In 2005, the Convention’s scientific advisory body adopted the LPI metric as part of a suite of biodiversity indicators, deployed to monitor progress towards that target ^25^. In 2010, the CBD Parties agreed a further set of biodiversity targets, the Aichi Targets, for the period 2011 to 2020 ^4^ and the LPI was identified as an indicator for several of these.

To strengthen the LPI’s scientific foundations and improve its capacity as an indicator for tracking progress towards international biodiversity policy targets, an in-depth peer-reviewed paper on the methodology was published ^18^ and the current partnership between WWF and the Zoological Society of London (ZSL) was subsequently formed in 2006. Since then, two updates to the methodology behind the global index have been published ^26,27^ and the research potential of the LPI data has expanded by incorporating metadata on ecology, geography, threats and management into the database, the core data of which were made openly accessible online in 2013 (18% of the data set is not available due to a confidentiality clause in the data sharing agreement, often for rare or threatened species).

### Applications of the LPI

Here we provide an overview of the uses of the different LPI project elements (see Box 1) and outputs, grouped into three themes: public engagement and advocacy, policy and research.

1. *The LPI as a communication tool for public engagement and advocacy*. From its inception, the LPI was seen as a powerful tool and WWF communications found that it resonated with the public better than any other conservation messages at that time. The LPI helps to set the scene for the state of global biodiversity by conveying a complex topic as a singular takeaway message for a broad audience. The key conduit for the global LPI has been as the headline biodiversity indicator within the (LPR). The LPR is an open access, biennial publication of the latest research and insights into global biodiversity trends, the human drivers behind them, and proposed solutions to halt biodiversity loss and “bend the curve” ^10^ back towards restoration. Its widespread distribution and WWF’s communications expertise have provided a regular global media platform and, emphasizing opportunities for awareness raising and advocacy regarding the biodiversity crisis. The 13^th^ edition, published in 2020, was translated into 16 languages and circulated around the world, with over 290 million social media views and 3,560 mentions from monitored global news outlets within the first month of its launch ^28^. The consistent use and media exposure within the LPR has accorded the LPI with familiarity within the public realm (see Communication and interpretation of the LPI). An analysis of online posts and articles (in English) containing the LPR 2020’s keywords or hashtags showed that 51% mentioned the 2020 global LPI statistic ^28^. Apart from global LPI figures, analysis of subset indices such as those featured in the LPR 2020 (LPI by The Intergovernmental Science-Policy Platform on Biodiversity and Ecosystem Services (IPBES) regions, taxonomic focus (e.g. reptiles) and ecological biome (e.g. forests and freshwater)) have been used to draw focus towards trends within different species groups ^5,29,30^. Both the underlying data in the LPI and the global results have been used in several educational formats, in schools and higher education. As part of the latest LPR outreach campaign, a youth edition including the LPI trends was prepared ^31^ and adapted by WWF country offices to enable young people to learn from the report’s key messages and promote engagement of schools globally in biodiversity issues. Nature documentaries provide another medium for large-scale biodiversity outreach ^32^. The 2019 Netflix series “Our Planet,” narrated by Sir David Attenborough, used the global LPI statistic from LPR 2018 to set the scene for its narrative alongside other headline biodiversity indicators and, within the first month of the launch, was viewed by 45 million accounts across the world ^33^. National scale LPI analysis and LPRs such as those undertaken by WWF offices in Belgium ^34^, the Netherlands ^35^ and Canada ^36^, and regional approaches like the 2013 “Wildlife Comeback in Europe” report ^37^ have used LPI figures to illustrate species trends and raise public awareness to what is happening to status and trends of the biodiversity on their doorstep. The Wildlife Comeback report reached 138 million people across Europe and worldwide ^38^.
2. *The use of the LPI project within policy* Analyses of the LPI dataset and trends within a geopolitical, ecological or taxonomic focus have been used to provide evidence of biodiversity change for policy makers, fed into policy and target development, and monitored progress towards those targets. The LPI is part of a suite of biodiversity indicators adopted by the CBD, measuring trends in relative abundance of vertebrates and deployed to monitor progress towards the 2010 Biodiversity Target ^19^, subsequent 2020 Aichi targets ^4^, and is one of the indicators within Goal A of the Post-2020 Biodiversity Monitoring Framework ^39^. As a measure of population trends compiled at annual intervals, the LPI is sensitive enough to detect annual changes, which is of value for informing policy ^15^ and evaluating the impact of conservation interventions ^40^.^19^ ZSL and WWF joined the Biodiversity Indicators Partnership (BIP) in 2007 to further develop the LPI and make it available for use under the CBD strategic plan. This resulted in the use of the LPI as evidence of biodiversity decline in international policy documents (Table 1): global and regional assessments (Millennium Ecosystem Assessment (2005) ^41^, IPBES global, regional and thematic assessments ^7,42-45^ and successive updates of UN Global Environment Outlook ^25,46-49^ and UN Global Biodiversity Outlook ^50-53^) as well as thematic assessments (Ramsar Convention on Wetlands, (2018) ^54^, Mediterranean Wetlands Outlooks (2012 and 2018) ^55,56^, the Convention on Migratory Species reports (CMS) (2008 and in 2019) ^57,58^ and Arctic Biodiversity Assessment (2013) ^59^). More recently, the global and regional indices were used to illustrate the state of nature and how this varies geographically as part of the evidence base for the Dasgupta review, an independent report on the economics of biodiversity ^60^.

**Table 1.**
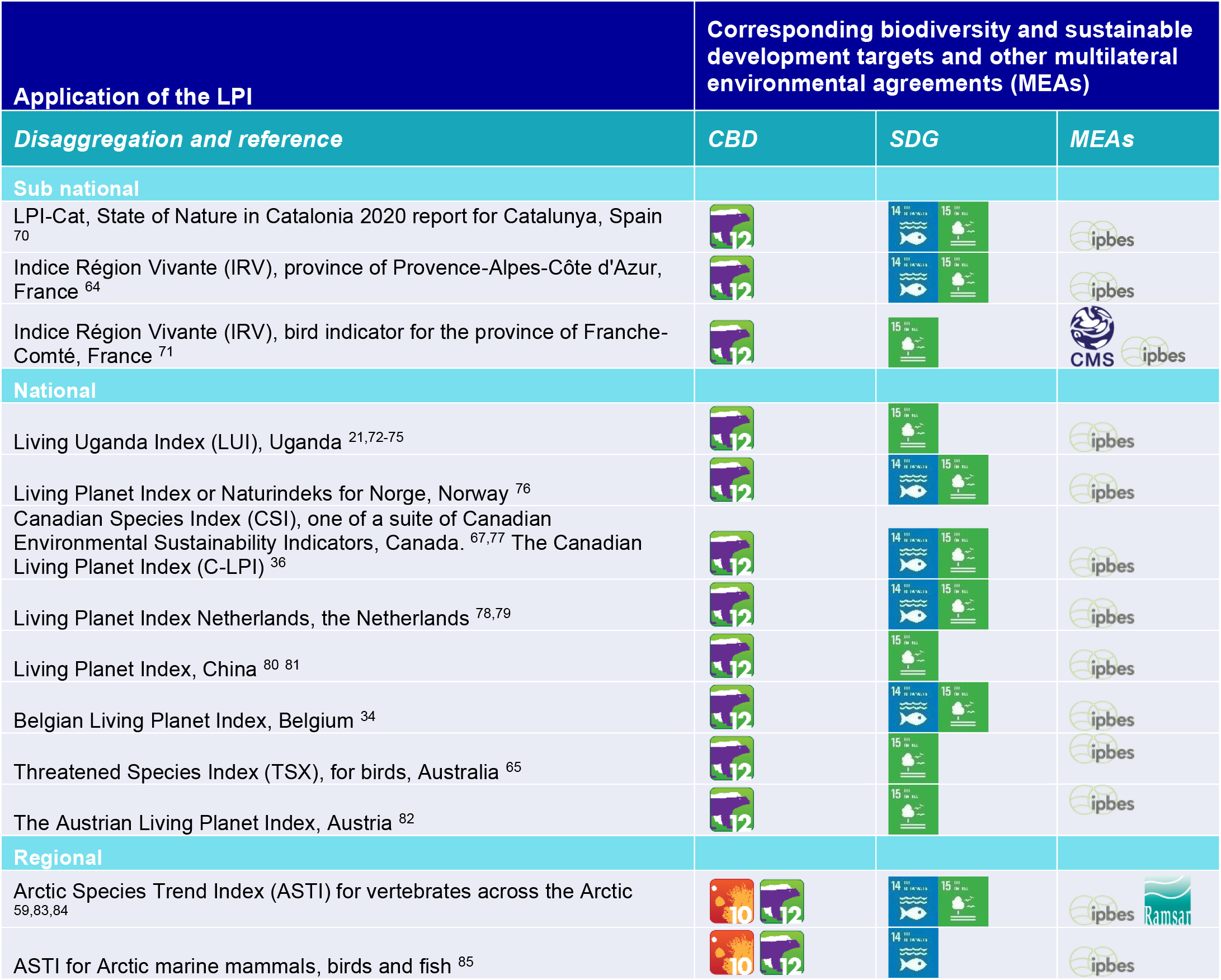
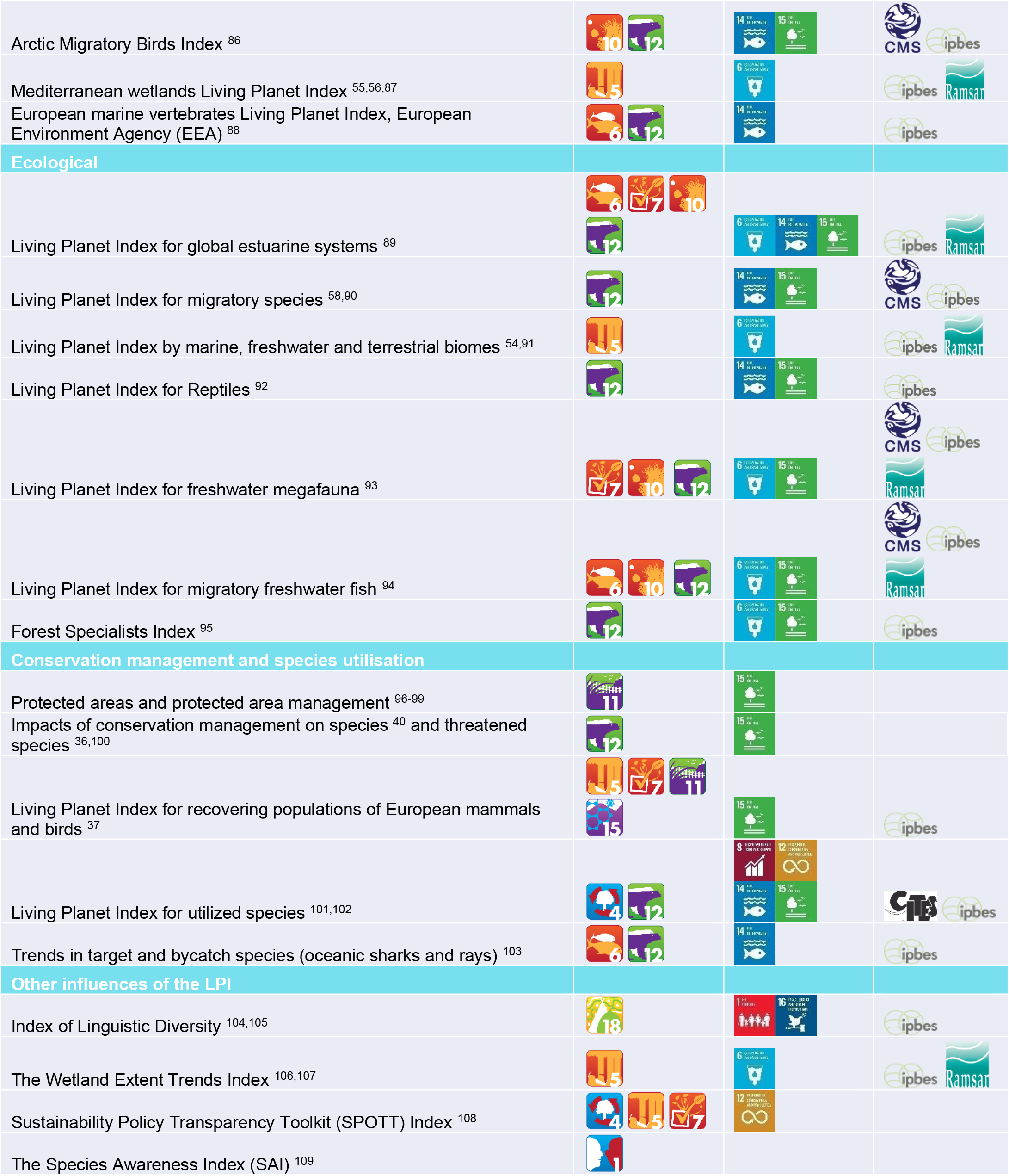
Selected applications of the LPI data and/or method and the corresponding and suggested uses for tracking global conventions on biodiversity, sustainable development, and other multilateral environmental agreements (MEAs). Sourced from: UNEP (United Nations Environment Programme) ^12,^UN ^13,^UNEP-WCMC (UN Environment Programme World Conservation Monitoring Centre) ^110^. The Post-2020 Biodiversity Framework targets were not finalised at the point of submission and are not included. LPIs have been used as a scientific basis and in their scene setting capacity, to influence policy development when advocating for transformative change and setting ambitious biodiversity targets ^9,10^. The global LPI statistic has featured in high-level biodiversity discussions, for example within Volkan Bozkir’s (President of the UN General Assembly) speech to heads of state at the 75^th^ UN Summit on Biodiversity in 2020 and within UK parliament in 2016 to support an Early Day Motion on Global Biodiversity ^61^. The LPI dataset and guidance on applying the method at a sub-global scale ^62^ have allowed for regional, national and in some areas, sub-national scale analysis (Table 1). This ‘scalability’ is a key requirement for indicators to be effective at tracking progress of signatory parties towards larger intergovernmental targets ^62,63^. CBD parties, for example, can develop national LPIs to fulfil part of their progress reporting requirements within their National Biodiversity Strategy and Action Plans (NBSAP) ^3^. Several members, including the Netherlands, Uganda, Canada, and China have provided LPI analysis of species trends within their NBSAP reports. In France, this process has been scaled down even further and; provinces such as Provence-Alpes-Côte d’Azur have used LPI analysis to track progress towards their National Biodiversity Strategy ^64^. In Australia, a new application of the LPI method focussed on threatened species to monitor their national progress towards Aichi Target 12 (extinction prevented) ^65^. Aside from tracking CBD commitments, nations have adapted the LPI method and applied it to suit their state biodiversity indicator needs such as the “Canadian Species Index,” developed by ZSL in partnership with Environment and Climate Change Canada (ECCC) ^66,67^. The package in the programming language “R” for calculating the LPI (rlpi), is freely available via GitHub ^68^, and has been used by collaborators from around the world to produce their own regional and national indices e.g. national and scientific agencies within Brazil use it within a national bird and mammal monitoring programme ^69^.
3. *The LPI project as a tool for research* The LPI methods, dataset and metrics have been used either individually or in unison for numerous research projects around the world (Table 1 and Figure 1). Within a random sample of 341 citations containing the term “Living Planet Index,” 90% of author and document affiliation was classed as research (academic institution or university); of the outputs themselves, 53% were within academic journals (Supplementary Materials A and B).

**Fig. 1.**
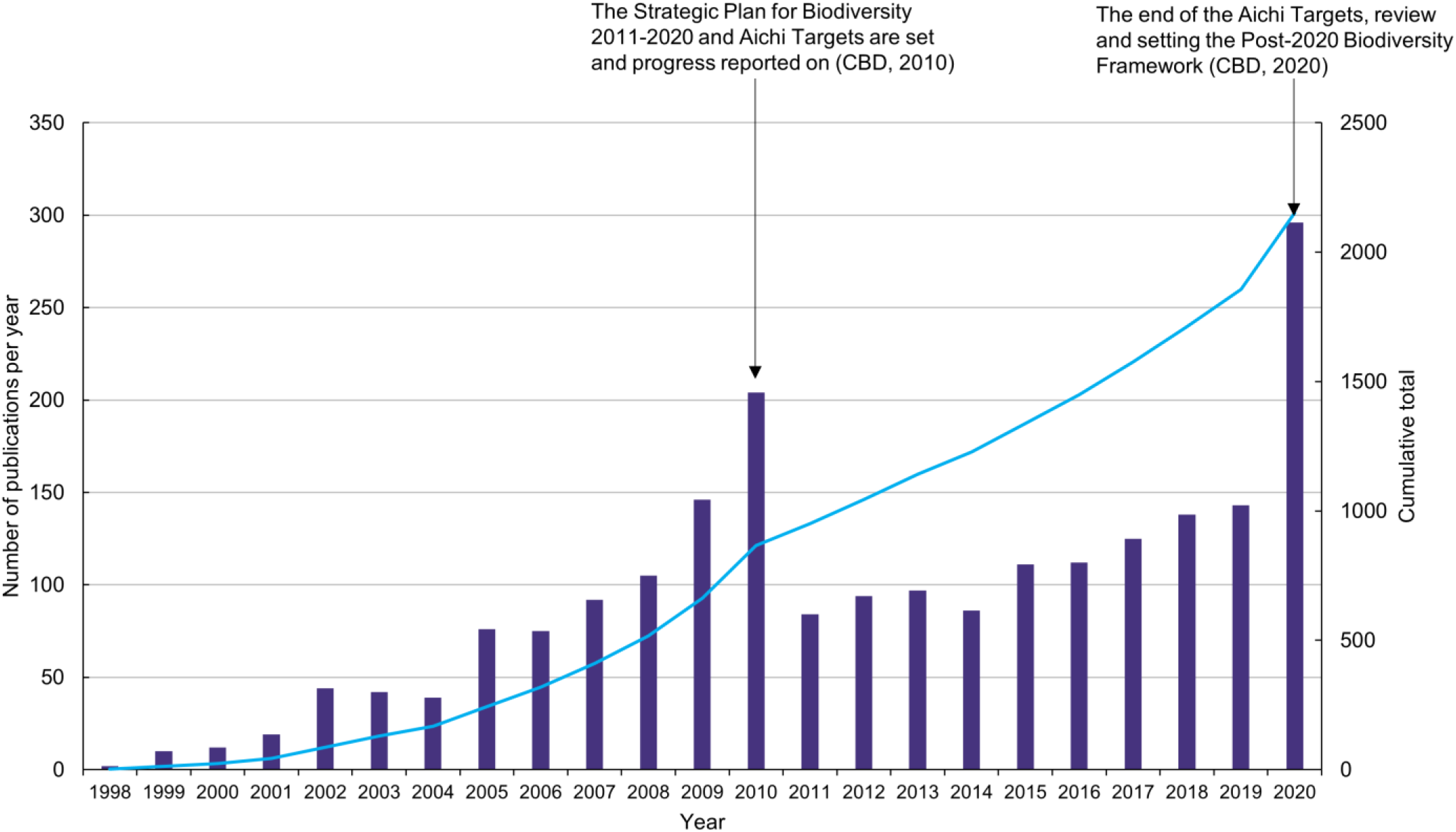
The number of publications per year citing the Living Planet Index between 1998-2020. The secondary Y-axis shows the cumulative total of publications. These 2,152 citations are from academic and grey literature in English and non-English languages between the years 1998-2020 (as of 18^th^ of January 2021). See Supplementary Materials A for details on the methods. The Living Planet Database (LPD) (except for about 18% of the data marked as confidential – see the origins and development of the Living Planet Index) has been publicly available since 2013 when the LPI website was created to facilitate viewing and downloading the data. Prior to this, subsets of the database were shared upon request. The LPD is now the largest repository of vertebrate population trend data (containing over 38,000 populations of more than 5,000 species at the time of writing), adding to a wealth of available biodiversity data for species occurrence (GBIF ^111^), species extinction risk (IUCN Red List ^112^) and ecological community data (PREDICTS ^113^, BioTIME ^114^). To date, www.livingplanetindex.org has had over 6,000 registered users from 145 countries around the world. Within the LPD, the population and ancillary data (Supplementary Materials D Figure 5) have facilitated a wide range of research topics (Table 1). In particular, the threat and management data at population-level allows for more fine-grained analysis compared with using species-level data. Recent applications of the data include: measuring the effectiveness of protected areas ^96-98^; evaluating the correlates of abundance trends in subsets of species such as mammals, reptiles, forest specialists, freshwater megafauna and migratory species ^90,92,93,95,115^; the nature of population dynamics in response to threats or management ^99-101,116,117^; the effects of land use and climate on species ^118^ and exploring linkages between human development variables and wildlife population trends ^119^. The LPD has been incorporated into an open access repository at the University of Edinburgh, dedicated to providing free online courses in statistics for ecology and environmental scientists ^120^. In a more informal setting, LPI data have been used to present challenges for data visualisation or analysis as part of Hackathons, one of which led to the development of a tool to automatically identify papers containing abundance data ^121^. The framework used to calculate the LPI has been applied to produce other metrics and not just for biodiversity. Conceptually, relative change, as calculated by the geometric mean, can be applied to other units of measurement that have been collected consistently over time. Using the code for calculating the LPI, new indicators have been developed for wetland areas ^106,107^, linguistic diversity ^104^, monitoring environmental, social and governance transparency in palm oil production ^108^ and biodiversity awareness ^109^. The first two of these are part of the ongoing suite of indicators for the CBD (Table 1).

### Challenges and opportunities

Along with other high-profile biodiversity indicators and reports ^122,123^, the underlying data, methods, and interpretation of the LPI have come repeatedly under scrutiny, which has been a positive catalyst for new research, collaborations and ameliorations on the scientific rigour of the index. Here we provide an outline of the challenges faced by the LPI and aim to provide clarity on common misconceptions that have arisen within recent years.

1) *The dataset underpinning the LPI* One of the strengths of the LPD is that it is not static: data are continually added and updated to provide the most complete and accurate picture possible of relative trends in population sizes (Figure 2). To ensure data are comparable, only species-level time-series which fulfil the following criteria are added: they are a measure of population abundance (or proxy, such as number of breeding pairs), with two or more years of data, collected within a specified geographic location under consistent methods (or explicitly corrected for) ^26^. Supplementary metadata (Supplementary Materials D) are continually updated for both new, and existing time-series, adding a further step in the data extraction process ^26^. The rigorous evaluation of data sources and data extraction not only limits the amount of applicable data that can be included, but it is also time consuming and labour intensive, and affects the volume of data that can be processed for each update. Storing these data in suitable infrastructure and the financial support required to maintain it are a further limitation common to other biodiversity databases ^124^. The costs of running the entire project can be complex to calculate as the source data are often already published and there are many stakeholders including researchers and policymakers to consider.

**Figure 2.**
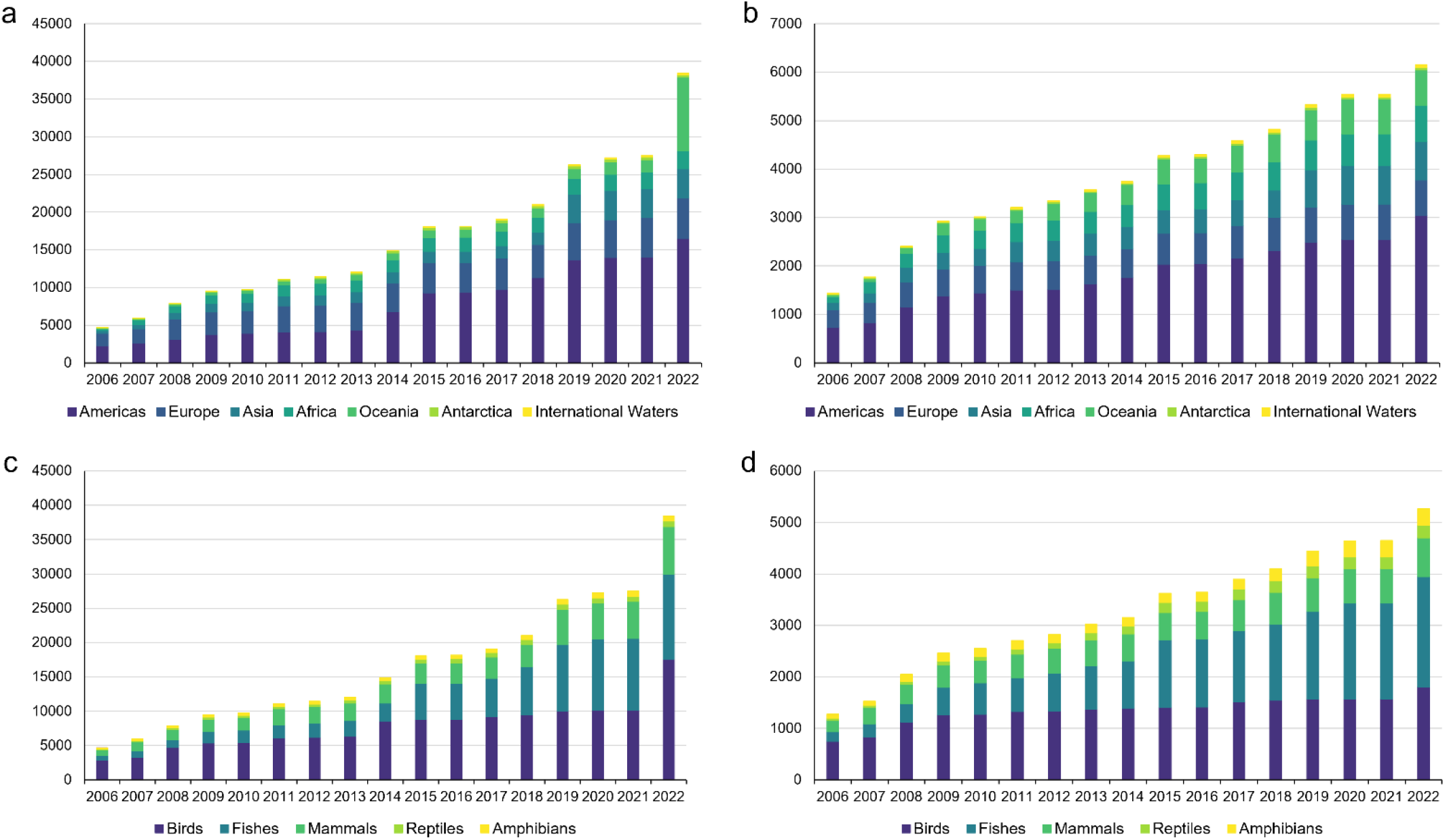
a-d. Growth in number of populations and species in the Living Planet Database (LPD) by region and taxa. The cumulative number of new populations (panel a) and species (b) entered by region, and the cumulative number of populations (c) and species (c) entered by taxon. Please note 2b adds up to more than the individual number of species as some species occur in more than one region. Long-term, population, abundance studies at a species population level are a limited resource in themselves, particularly for highly speciose taxa such as invertebrates and plants which have not been included in the LPD to date (see The future). Studies which include population data may not have been designed for long-term population monitoring but to assess population size and so their methods and survey effort might change with advances in population estimate approaches (e.g. revised Orangutan estimates in Sabah ^125^, this renders these data incompatible for inclusion in the LPD). This issue is amplified for regions and taxa which are recognised as underrepresented within the dataset such as tropical regions and fish, reptiles and amphibians (see Supplementary Materials E) ^27,126^. Subsequently, the composition of the LPD is likely to reflect bias inherent in species monitoring schemes which tend to favour certain taxa (e.g. birds), or regions (e.g. high income countries) ^27,127,128^. This is a challenge shared by biodiversity indicators and databases in general ^122,129^. In addition, attempts to source data from grey literature or offline databases is often dependent on the time and expertise available from researchers and field contacts within chronically neglected and underfunded areas ^130^. To counteract bias in the resulting LPI, two approaches are taken. At the data inputting stage, a gap analysis of the taxonomic and geographic representation of the LPD is used to prioritise taxa and regions for targeted data searches (Supplementary Materials E). However, focussed searches are not always fruitful: within the 2020 LPI, only 4 populations of African amphibians were included despite targeted efforts ^131^. The second step for overcoming bias in the LPI is in the adoption of the diversity-weighted method (see The LPI methods). Language is a further constraint to collating representative data for the LPI and can exacerbate existing geographic biases ^132^. The dominance of English-language data sources is partly a reflection of the LPI project being hosted in an English-speaking country but also of English as a globally used language for science ^133^. However, over a third of biodiversity documents from a single year were published in languages other than English ^134^, so there are likely to be data that have not been captured because language barriers have not yet been adequately addressed. Collating and storing a continually increasing repository of LPI data, that aligns with FAIR (Findable, Accessible, Interoperable, and Reusable) Data Principles, requires ongoing investment in the data infrastructure and management ^124,135^. Coupled with this is the importance of promoting data sharing in a way that alleviates concerns over data ownership and provides appropriate credit to data providers. Unless a system is in place whereby data providers maintain ownership and control of their data, there is likely to be a barrier to mobilising data.
2) *The LPI methods* The key methodological challenges for the LPI project are to generate a robust LPI indicator of biodiversity and to model the time-series data in the LPD, which vary in length and scale, in a way that allows exploration of underlying patterns in population trends. A further challenge that underpins both issues, is addressing the taxonomic and geographic gaps in the underlying data (Supplementary Materials E). The basic formula for calculating the LPI has remained consistent: each logged population trend is averaged within a single species and the species trends are aggregated to produce a single index ^18^. This aggregation is produced using a geometric mean, an approach used to generate other indices of relative abundance from species abundance data ^136-138^. Further levels of aggregation are often used for global, national, and local contexts (see Supplementary Materials F for the global example). A challenge in the use of a geometric mean of abundance for the calculation of indicators is that it can be sensitive to outliers in the data which may impact the precision of the long-term trend if not addressed ^139-141^. While this method is still considered to be a more suitable and sensitive metric to assess changes in biodiversity ^142,143^, understanding the impact of outliers is important. To tackle this, each new iteration of the global LPI analysis includes sensitivity tests on the influence of single species on the trends and of the effect of short time-series on the LPI, as these are more commonly associated with highly variable or extreme trends ^131^. These tests are published in the supplementary information, blog or website for transparency and to demonstrate the robustness of any index ^131^. The modelling of the time-series data in the LPD has been periodically improved. In early iterations of the LPI, the chain method was implemented, which involved linearly interpolating the rate of change between 5-year intervals, (following Loh, et al. ^18^). As this approach was sensitive to abrupt changes in population trends, generalised additive modelling (GAM) was adopted to better capture long-term nonlinear trends in populations ^26^. National variations of modelling have been tailored to the type of species monitoring data in the country in question, for example the use of linear regression for short-term trends in the Canadian Species Index ^66,79^. More recently, Bayesian approaches such as state-space models have been applied to model the population time-series whilst incorporating observation error into the estimation of trends ^144^, something that the GAM framework does not account for. This has allowed for new ways of analysing the LPD, which lend themselves to uncovering the correlates of vertebrate population trends ^145^ and the taxonomic and geographic patterns of population trends globally ^141^. A significant challenge remains in tackling the underrepresentation in the LPI database of particular taxa and regions in the LPD so an adaptation to the LPI method was made to mitigate the impacts of this bias on the index. This diversity-weighted approach was developed and subsequently adopted for calculating global and regionals LPIs ^27^. This method places greater weight on species trends from regions and taxa that are more species-rich but tend to be disproportionately under-represented in the LPD e.g. the Neotropics. This provides a more representative picture of global vertebrate trends in lieu of a more complete data set. One drawback is that weight is often placed on species and regions with the lowest data availability so if the sample of data from a region is not representative, this could cause an over-or underestimation of trends. Efforts are also underway to address gaps in the data set through targeted data collection and to develop models to predict trends in locations and for taxa which are data deficient, as has been done for extinction risk ^146^.
3) *Communication and interpretation of the LPI* Key attributes of biodiversity indicators are that they should be simplified and easily understood ^138^. The LPI was developed with these criteria in mind and, by aggregating trends from different ecological realms and geographic regions, it can provide a useful overview and communication tool for broad audiences. However, the index has been critiqued as oversimplifying the state of biodiversity ^147^ and masking important trends ^141^. Arguably, there is need for a balance between providing a simple, clear message about global biodiversity trends whilst supporting it with more in-depth analysis ^123^. To explore this variation, disaggregations of the LPI have been developed (Table 1), for example, for forest specialists ^95^. The limited availability of quality, ecological data prior to the 1970s is a common limitation to many biodiversity indicators ^129,148^. The LPI is benchmarked at a temporal baseline of 1970, and this raises the importance of interpreting the index in context, as geopolitical regions have been impacted by anthropogenic pressure at different points in time and varying intensity. In Europe, for example, a significant amount of habitat destruction and overexploitation of some species had occurred prior to the 1970s and therefore the LPI baseline is set at a significantly depleted reference point ^37^. The year chosen as a baseline can affect the interpretation of the state of biodiversity in a particular region ^149^. Without taking this into consideration, it is possible to underestimate the gravity of the decline in biodiversity or overestimate a recovery within any given landscape. Communications around biodiversity indicators and biodiversity loss have often centred on species, and species extinctions respectively, rather than attempting to explain the multi-faceted nature of biodiversity change and how we measure it ^123,150^. Miscommunication and oversimplification of biodiversity and biodiversity loss, or decline, across the science-society and science-policy interface, are a challenge shared by biodiversity indicators in general ^123^. The impact of substituting a single word for another in press and media communications, “loss” vs “decline”, has sometimes led to misinterpretation of the global LPI statistic. A negative trend in the LPI depicts a relative decline in population sizes, on average, since 1970. The use of the word “loss” in some media articles can imply that a negative LPI trend is analogous with the disappearance of populations and even the extinctions of species, which can prove challenging to correct. Media headlines have referred to large percentages of populations being “wiped out” ^151^, which could mislead the public about the severity of biodiversity decline and, it has been argued, such negative statements about environmental issues may be counterproductive in trying to stimulate action ^152^. Efforts to minimise misinterpretation have been made with each iteration of the LPI, by engaging with journalists directly through press briefings, providing background information to communications teams and publicising the supporting information available in technical supplements to the LPR ^131^, websites (http://stats.livingplanetindex.org/) and blogs ^153^. These efforts have also reinforced the LPI as a measure of “relative abundance” rather than “abundance” to help avoid misinterpretations ^154^. We have already seen an uptake in the use of the LPR 2020 technical supplement in recent publications and blogs exploring the LPI ^155,156^. The analogy of a FTSE index for biodiversity is most commonly used to describe the LPI, but a focus in the future should be on finding other ways to communicate the index that mitigate the use of dramatic narratives, whilst retaining the simple message of the LPI that can be broadly understood.

### The future

The LPI project has grown significantly over the last 25 years and provides an important dataset to communicate the trends in vertebrate populations and investigate the factors that influence them. We identify four key priorities for the immediate future.

#### Increasing representation in the LPI

The composition of the LPI needs to be improved, crucially by increasing the taxonomic and geographic representation of the data particularly for aquatic species. Incorporating invertebrate and plant species into the LPI is likely to be challenging given the paucity of monitoring compared to some vertebrate groups ^157^ but is key to attaining an indicator of broader biodiversity in addition to and providing a fuller data set for macro-ecological research. Many national LPIs have already been developed, and maintaining this focus on increasing the representation of species within countries will provide nations with a tool to track progress towards future CBD and SDG targets. Indicators also need to be ecologically relevant ^138^, so ensuring that different functional attributes of species within an ecosystem are reflected will be the focus of new research. These developments in the data set will be realised through the use of emerging techniques to incorporate unstructured data, such as that collected through citizen science initiatives ^158-160^, and capitalising on growing technology for monitoring biodiversity such as eDNA, satellite monitoring and AI-assisted counting of species, provided they can be transformed into usable metrics of abundance.

#### Streamlining data collation and data access

Sourcing and extracting data continue to be significant bottlenecks for the development of the LPD. Data searches can be automated to some degree using predictive models based upon titles and abstracts ^121^, but extracting data automatically remains a challenge. Working with publishers, data holders, government institutions and research funding bodies to automate the process of identifying and extracting data from articles would be beneficial particularly if a standardised workflow is developed (e.g. Cardoso, et al. ^161^), and systematic review tools may advance data collation in a community-driven way ^162^. To address language barriers, which in turn could help to fill taxonomic and regional data gaps ^163^, a protocol for conducting data searches in multiple languages is under development. This should be part of a broader strategy to build a sustainable data network for the LPI, which provides accessibility to a global database (both for data download and upload, e.g. from new national LPI datasets) whilst retaining data quality and ownership, and assuring appropriate credit to data gatherers and providers. It is also important that the LPD is made as accessible as possible, both through simple, downloadable, tidy data formats ^164^ and the development of Application Programming Interfaces (API) to allow the data to interoperate with other resources such as the IUCN Red List ^112^, Protected Planet ^165^ and GBIF ^111^.

#### Better models to link population trends with drivers

The LPI continues to highlight that global biodiversity is in trouble and understanding (and predicting) which regions and species are likely to decline most in the future is useful. As such, models to better predict wildlife abundance trends for species and regions where we have poorer data is critical. Understanding the quality and utility of these models will allow us to make concrete and valuable predictions. The varied response of some populations to their changing environment highlights an important question – are some populations useful ‘canaries’ of pending ecosystem collapse and how might we best identify them?

Models that combine LPI data with drivers such as land-use and climate-change data have demonstrated that both are important drivers of population trends ^118^. Developing these models further allows us to make predictions about how biodiversity might change under future scenarios and management interventions ^9^, highlighting one evolving use of biodiversity datasets like the LPD.

Whilst incorporating data on drivers from other global data sets can inform explanatory analysis for species trend data ^118^, population-scale information can also provide a powerful set of variables, for example in understanding the effect of different direct drivers ^101^ or to pave the way for counterfactual analysis of different management types (e.g. Jellesmark, et al. ^166^). However, the current coding for threats and conservation action in the LPD lacks alignment with established frameworks ^167^, so transferring the ancillary information into these classification schemes and maintaining the recording of population drivers will improve the utility of models and ground-truthing of broad scale datasets in the future.

#### Increasing the utility of the LPI for policy

From a policy perspective, an emphasis on developing LPIs at the national level is needed to expand its use as a communication and reporting tool. With reporting requirements at a national level for the SDGs and the CBD, national LPIs would serve a dual purpose of providing countries with a sensitive indicator for reporting while boosting data representation for the global index. Disaggregations of the LPI on themes such as use, trade, migration and wetlands should continue to be developed, so that these are available for reporting against other multilateral environmental agreements such as the Ramsar Convention on Wetlands, CITES and the CMS.

The LPI performed well in an evaluation of biodiversity indicators using decision science ^17^, although gaps were identified in the practice of regular tests of the index and in assessing the cost-effectiveness of the LPI relative to other indicators. Creating a better understanding of how the LPI fits within the growing suite of biodiversity indicators such as the Red List Index ^168^ and the Biodiversity Intactness Index ^169^, and clearly presenting the complementarity of these indicators with each other, will be key to developing a clear and consistent narrative of global biodiversity change ^14^ and to ensure the suitability of the LPI within any multi-dimensional indicator framework ^170,171^.

## Conclusion

The LPI has evolved from a simple communications tool to a large and growing database, policy tool and foundation for research. The open-access dataset and method are globally important resources for the scientific community and beyond, but improvements are still needed to enhance the representation of biodiversity in the underlying data and produce clear and meaningful outputs. Collaboration and engagement within the fields of science, policy, conservation and communication — some of which have fuelled much of the development to date, will continue to be important for ensuring the LPI project remains fit for purpose.

## Acknowledgements

This research was partly funded by Research England, SL was funded by WWF-NL; LM, SD, VM were funded by WWF-UK.

We acknowledge the following individuals who were instrumental in the initial development and/or funding of the LPI project; Georgina Mace, Ben Collen, Jonathan Baillie, and Raj Amin. We also thank the LPI volunteers, collaborators, and contributors to the LPD past, and present for their essential support to the LPI project.

## Supporting information

Supplementary information is available for this paper at: DOI XXX

These include the methods for the citation search in academic and grey literature, and metadata coding (Supplementary materials A), the results of citation search and metadata coding (Supplementary materials B), the results of the Altmetric data analysis for key LPI papers (Supplementary materials C), an infographic of the underlying data within the LPD and summary of the growth in populations and species in the LPD over time (Supplementary materials D), a summary of LPD data diagnostics for underrepresented taxa and realms (Supplementary materials E) and a visualisation of the global aggregation LPI method (Supplementary materials F).

