## Supplementary materials for "Past, present, and future of the Living Planet Index"

[1] Institute of Zoology, Zoological Society of London (ZSL), London, UK
[2] School of Anthropology and Conservation, University of Kent, Canterbury, UK
[3] WWF Netherlands – World Wide Fund for Nature, Zeist, Netherlands
[4] Global Center for Species Survival, Indianapolis Zoo, Indianapolis, USA
[5] School of Biological Sciences, University of Bristol, Bristol, UK
[6] WWF Canada – World Wildlife Fund, Toronto, Canada
[7] Institut de recherche pour la conservation des zones humides méditerranéennes, Tour du Valat, Arles, France
[8] Environment and Climate Change Canada (ECCC), Government of Canada, Gatineau, Canada
[9] WWF UK – World Wide Fund for Nature, Cambridge, UK
[10] Conservation and Policy, Zoological Society of London (ZSL), London, UK

#### Methods for citation search in academic and grey literature, and metadata coding

##### *Citation search and data collection*

Three online platforms of scholarly literature were selected for their complementary coverage of both academic and grey literature and to boost representation of non-English language texts which are often neglected from citation search tools [1]; Scopus, Web of Science and Google Scholar (via the academic citation analytical program Publish or Perish). Keywords chosen for the search were kept as the 'Living Planet Index' across all platforms for consistency and the acronym 'LPI' was not included to reduce the number non-target citations. The timeframe for the search was limited to 1998-2020 to coincide with the development of the LPI and to avoid non-target results from literature prior to the LPI's existence. We note that excluding searches for variations of the 'Living Planet Index' in English and other languages will potentially reduce the representation of LPI use at national and subset levels.

- **Data screening.** The results of the three citation search platforms were combined into a singular database and screened for search return errors, duplicate entries, Living Planet Report publications and incomplete entries, which were removed from further analysis. Publications with authors from the Indicators and Assessments Unit were coded so they could be differentiated from external users in the future.

- **Language classification and screening.** The language for each entry was classified from International Organization for Standardization (ISO) language codes using the text within the publication title and two formulas within Google Sheets (ISO language detect and Google Translate functions). Language classifications were screened and erroneous allocations (misinterpreting author names, years, abbreviations and Latin) were corrected.

##### *Random sample metadata coding*

To explore patterns in usage and reach in more detail, a random sample of 341 English language texts were coded with metadata such as geographical scale of focus, how the source used the LPI, and source type (See supplementary Table 1). Originally, we aimed to use the excerpt of text which Publish or Perish extracts from each source publication from Google Scholar to categorise the use of the 'Living Planet Index' for that entry. However, these text 'snippets' were limited in character length, not available for every source and would only return the 1st time the search term was used within the source. To evaluate the use of the LPI within a publication it was determined that returning to the source publication would be necessary.

- **Random sample**. Given time limitations and time taken to access each source publication and interpret the text and code up fields of interest (informed by a time trial of coding 50 random entries) it was determined that a random sample of >320 English language entries would be selected to act as a representative sample for a week's worth of coding.

- **Coding the sample**. Fields selected to be manually coded from interpreting the source document were categorising the use of the LPI within the document, the type of document, document affiliation, whether the focus of the document was at a global scale or a focal country, whether the document was accessible online, and a notes field. Definitions for LPI use, document type and document affiliation can be found in Supplementary Table A 1. To keep the source document focal country field standardised and to avoid typos, a data validation tool was used with the IPBES country list. The associated IPBES region was then automated using VLOOKUP.

- **Data screening**. We recorded whether the source document was irretrievable, or the citation had been misattributed by citation search tool error or by author error and removed these entries from analysis.

| Use of the LPI | | Document type | | Source affiliation | |
|---|---|---|---|---|---|
| **Category** | **Definition** | **Category** | **Definition** | **Category** | **Definition** |
| **Discussed** | The LPI is discussed in detail | **Journal article** | Published in a peer reviewed journal - to include data paper, editorial and review | **GOV** | Government or State; agency, commission, department, ministry, administration - government controlled or linked organisation |
| **Figures cited** | Figures such as percentages for different data cuts, species numbers etc are presented in some detail. | **Research outputs** | Preprint academic article, Conference papers published in the context of an academic conference or workshop and Doctoral dissertations/Theses | **NGO** | Non-governmental organisation - independent of government - usually not for profit |
| **Mentioned** | The LPI is referred to by name. This can be a very brief mention, e.g., in a list of indicators, or a slightly more detailed one which does not present figures from the LPRs. | **Books and book chapters** | Books and book chapters | **IGO** | Intergovernmental organisation/ international organisation - made from sovereign states (referred to as member states), or of other intergovernmental organisations |
| **Method used** | The LPI method is applied to other data | **Policy statement** | Specify which type in Author/Affiliation field | **PRI** | Private sector - for-profit businesses that are not owned or operated by the government |
| **Data used** | Data from the LPD is used but analysed using a different method | **Report** | Gov report, Stats report, Research report. Important to specify which Author/Affiliation type in that field | **RES** | Research institutes and centres, university, and academic institutes |
| **Method & data used** | The LPI method is applied to data from the LPD | **News or Media article** | Published online on a news agency website, magazine, or organisations website | **MISC** | Miscellaneous - make a note about it in the notes field |
| **Not named** | Any of the main publications (Loh paper, Collen paper, LPRs) are cited but the LPI is not used by name. | **Teaching material** | Presentations, syllabus, Guides for Classroom, teacher, lecturer, Non-classroom, documents for education purposes | **MULTI** | Multidisciplinary - a combination of different sectors as authorship |
| **Does not appear** | The LPI/ Living planet Index does not appear in the text - erroneous retrieval from search | **Other** | Not a research output - could be; Surveys, Letters, Blog posts, Newsletters etc - make a note in the notes field | **MEDIA** | Journalists, News agencies |
| **Error in use** | The LPI is named but the authors are referring to something else -e.g., footprint analysis or LPR | | | | |
| **Irretrievable** | Cannot get access to the document to assess LPI use | | | | |

**Supplementary Table 1. Definitions for metadata coding of the citation search results by usage of the LPI, document type and document affiliation**

### A. Results of citation search and metadata coding

The Living Planet Index was cited by a total of 2,152 academic and grey literature documents in 32 languages (including English) between 1998-2020 (see Supplementary Table 2 and Supplementary Figure 1).

Of the total 1,805 English language citations, a random sample of 375 entries were coded with metadata (21% of total); of these entries, 34 were excluded (9% of the random sample) due to the source being irretrievable or due to search tool or author error in attribution (see Supplementary Tables 2, 3 and 4), thus leaving a random sample of 341 entries.

| Citation search tool | Number of publications containing the phrase 'Living Planet Index' 1998-2020 |
|---|---|
| Web of Science | 46 |
| Google Scholar via Publish or Perish | 2,171 |
| Scopus | 526 |
| Total unique entries* | **2,152** |
| Total random sample of English language entries with complete metadata | **375** |

**Supplementary Table 2. Results of the citation search for 'Living Planet Index' within literature between 1998-2020.** *Unique entries refer those which have been through a data screening and cleaning process to remove duplicate entries, entries pre-1998, entries with no year and entries which contained the "Living Planet Report" in their title.

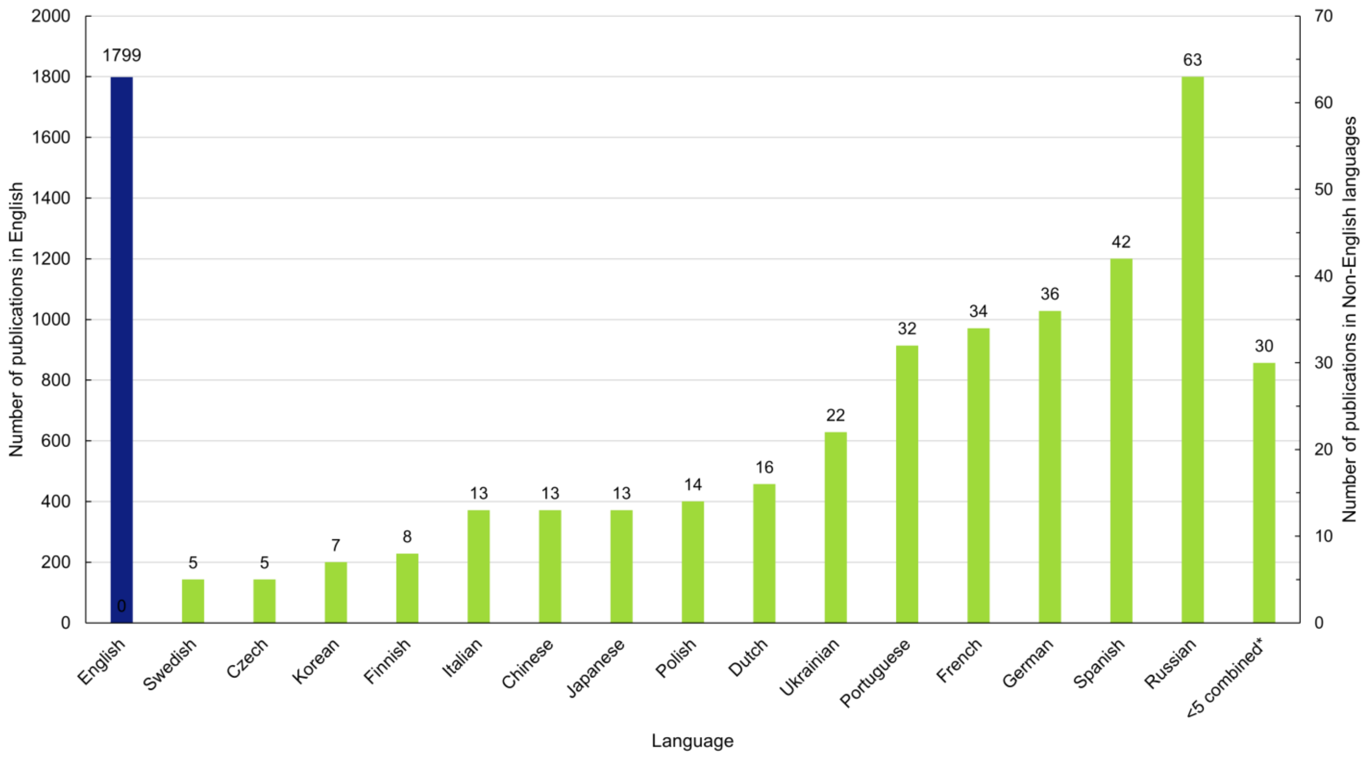


**Supplementary Fig. 1. Number of publications citing the Living Planet Index within English (left hand Y-axis) and non-English language (right hand Y-axis) texts between 1998-2020.** It is likely that LPI use in non-English languages will have been underestimated despite using the Publish or Perish software. * *The following 16 languages had <5 publications; Greek, Catalan, Croatian, Slovenian, Slovak, Lithuanian, Thai, Estonian, Galician, Bosnian, Danish, Romanian, Norwegian, Chinese (Traditional), Hungarian and Indonesian*

Metadata analysis of what aspects of the LPI project are being used, and how, revealed the most common uses were: mentioning the LPI (31%) and citing LPI figures (30%) with the majority of citing documents focused on the global scale (70%) (See Supplementary Materials B, Figure 2).

| Use of the LPI with document | Number of citations | Percentage % |
|---|---|---|
| Mentioned | 120 | 35% |
| Figures cited | 117 | 34% |
| Not named | 47 | 14% |
| Discussed | 23 | 7% |
| Method & data used | 15 | 4% |
| Method used | 12 | 4% |
| Data used | 6 | 2% |
| **Total** | **341** | **100%** |

**Supplementary Table 3. Results of the coding of how the LPI was being used with a random sample of English-language.**

Ninety percent of author and document affiliations were classed as research (academic institution or university, see Supplementary Materials Table 4); of the outputs themselves, 52% were within academic journals (see Supplementary Materials B Table 5).

| Author type or affiliation | Number of citations | Percentage % |
|---|---|---|
| Research | 306 | 90% |
| NGO | 10 | 3% |
| IGO | 10 | 3% |
| Governmental | 8 | 2% |
| Media | 4 | 1% |
| Private sector | 3 | 1% |
| **Total** | **341** | **100%** |

**Supplementary Table 4. Results of the coding the affiliation of authors and documents which cite the LPI within a random sample of English-language. Definitions of how the affiliations were classified can be found within in Supplementary materials A Table 1,**

| Source | Document type | Number of citations | Percentage % |
|---|---|---|---|
| Academic literature | Journal article | 179 | 52% |
| Grey literature | Books and book chapters | 69 | 20% |
| | Research outputs | 53 | 16% |
| | Report | 29 | 9% |
| | News or Media article | 5 | 1% |
| | Teaching material | 3 | 1% |
| | Other | 3 | 1% |
| **Total** | | **341** | **100%** |

**Supplementary Table 5. Results of the coding the source type and document types which cite the LPI within a random sample of English-language.**

Within regional (9%) and single country focussed documents (21%), the distribution by IPBES region was slightly biased towards Europe and Central Asia (36%) and Asia Pacific (26%) but all regions were represented (See Supplementary Materials B Figure 3).

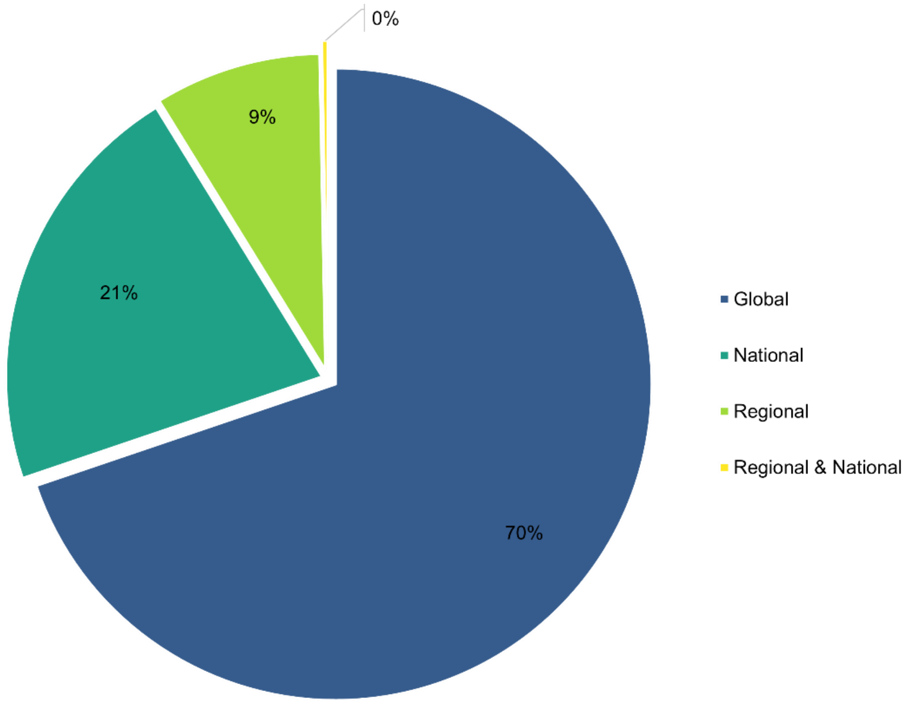


**Supplementary Fig. 2. Geographical scale of English-language publications which cite Living Planet Index (%).** A random sample of 341 English language publications were categorised as having a Global, National, or Regional scale focus.

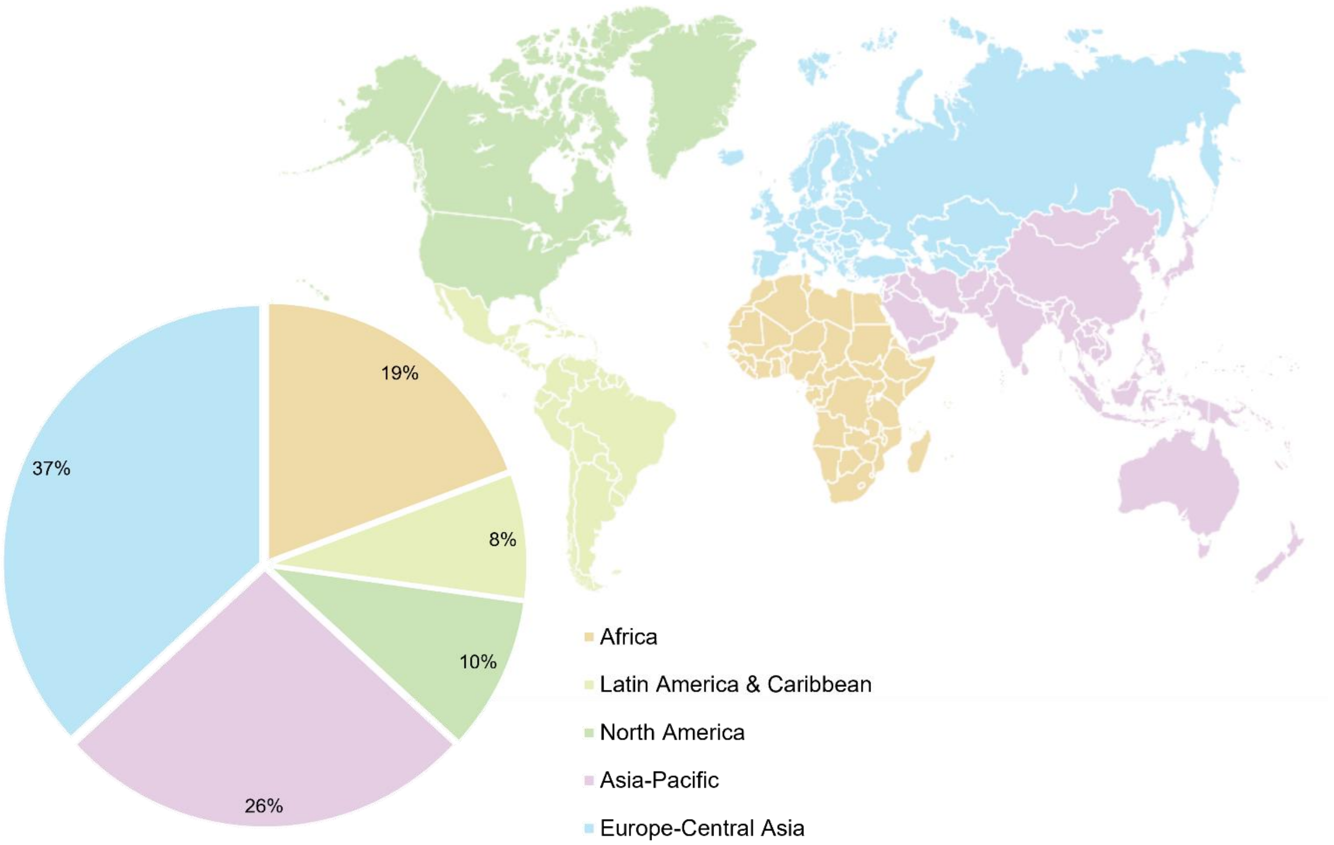


**Supplementary Fig. 3. Geographical distribution of national and regional focussed English-language sources which cite the Living Planet Index, by IPBES regions (%).** A random sample of 341 English language sources were categorised as having a Global, National, or Regional focus; of these 103 entries of National and Regional level sources were identified and attributed to IPBES regions. Within the national coded data, 37 individual countries were recorded (when counting Wales and England with the UK). Sources marked as multiple IPBES regions refer to the following: Arctic, Neotropics, Mediterranean and a study that looked at trends in both Canada and Sub-Saharan Africa. Map adapted from WWF [2] and originally sourced from IPBES [3].

#### B. Results of Altmetric data analysis for key LPI papers

As a means of inferring outreach within the scientific community, Altmetric analysis was undertaken on three key academic papers on the LPI method [4-6] and two CBD target reviews which included the LPI [7,8]; this revealed that these five papers were in the top 5% of all research outputs scored by Altmetric and had high attention scores ≥ 95 percentile (%) compared to outputs of the same age (See Supplementary Materials C Table 6 and Figure 4).

| Focus | Publication | Attention score | In top percentage of all research outputs scored (%) | High attention score compared to outputs of same age (%) | High attention score compared to outputs of the same age and source (%) |
|---|---|---|---|---|---|
| The Living Planet Index method | Loh, J. *et al* 2005 | 45 | 5 | 95 | 81 |
| | Collen, B. *et al* 2009 | 40 | 5 | 96 | 95 |
| | McRae, L. *et al* 2017 | 130 | 5 | 99 | 94 |
| Using the Living Planet Index and reviewing progress towards policy targets | Butchart, S. *et al* 2010 | 113 | 5 | 98 | 98 |
| | Tittensor, D. *et al* 2014 | 253 | 5 | 98 | 90 |

**Supplementary Table 6. Altmetric impact analysis of three key Living Planet Index methodology publications [4-6] and two policy target review papers [7,8]**. Data correct at the time of collection on 19th February 2021.

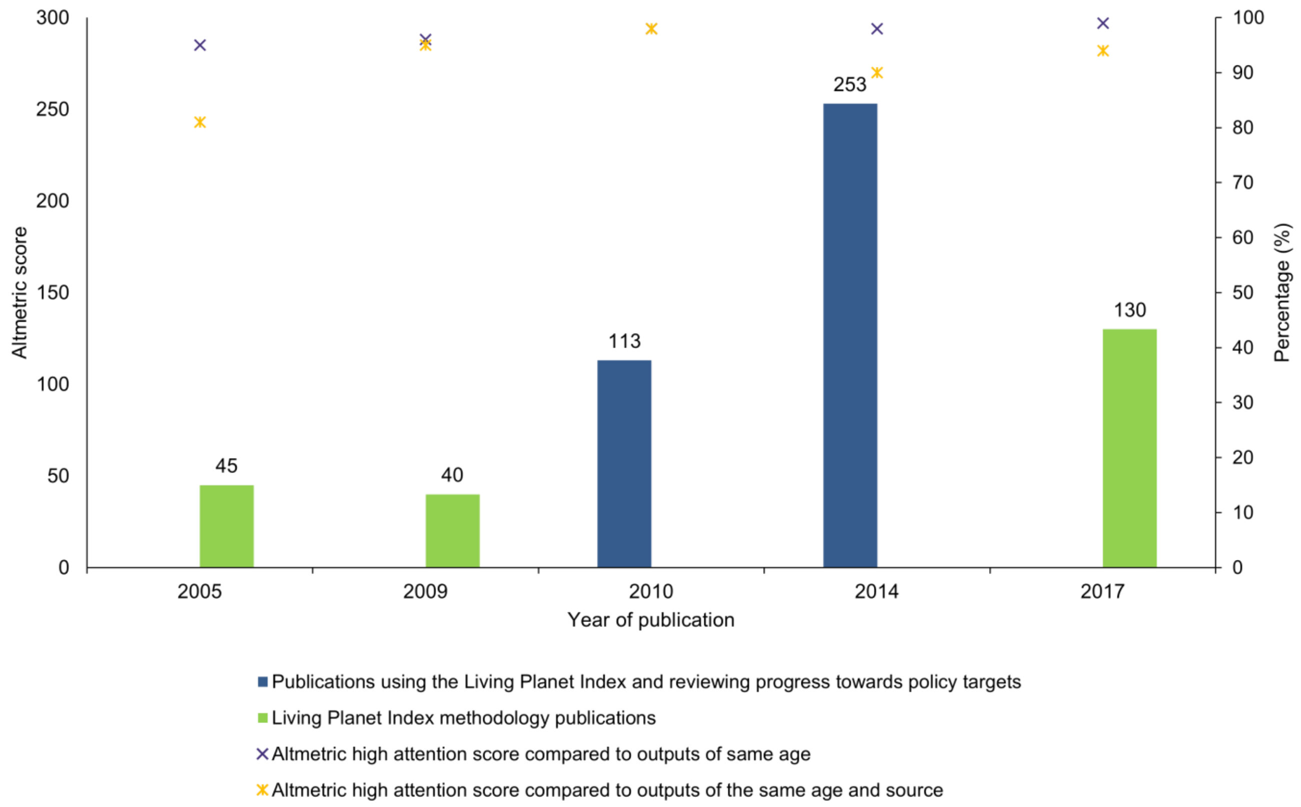


**Supplementary Fig. 4. Altmetric scores of the three key Living Planet Index methodology publications [4-6] and two policy target review papers [7,8] on the primary Y-axis.** The percentage of high attention scores compared to outputs of the same age, and same age and source are plotted on the secondary Y-axis. All publications were classed as within the top 5% of all research outputs scored by Altmetric. Data correct at the time of collection on 19th February 2021.

#### C. A visualisation of the underlying data within the LPD

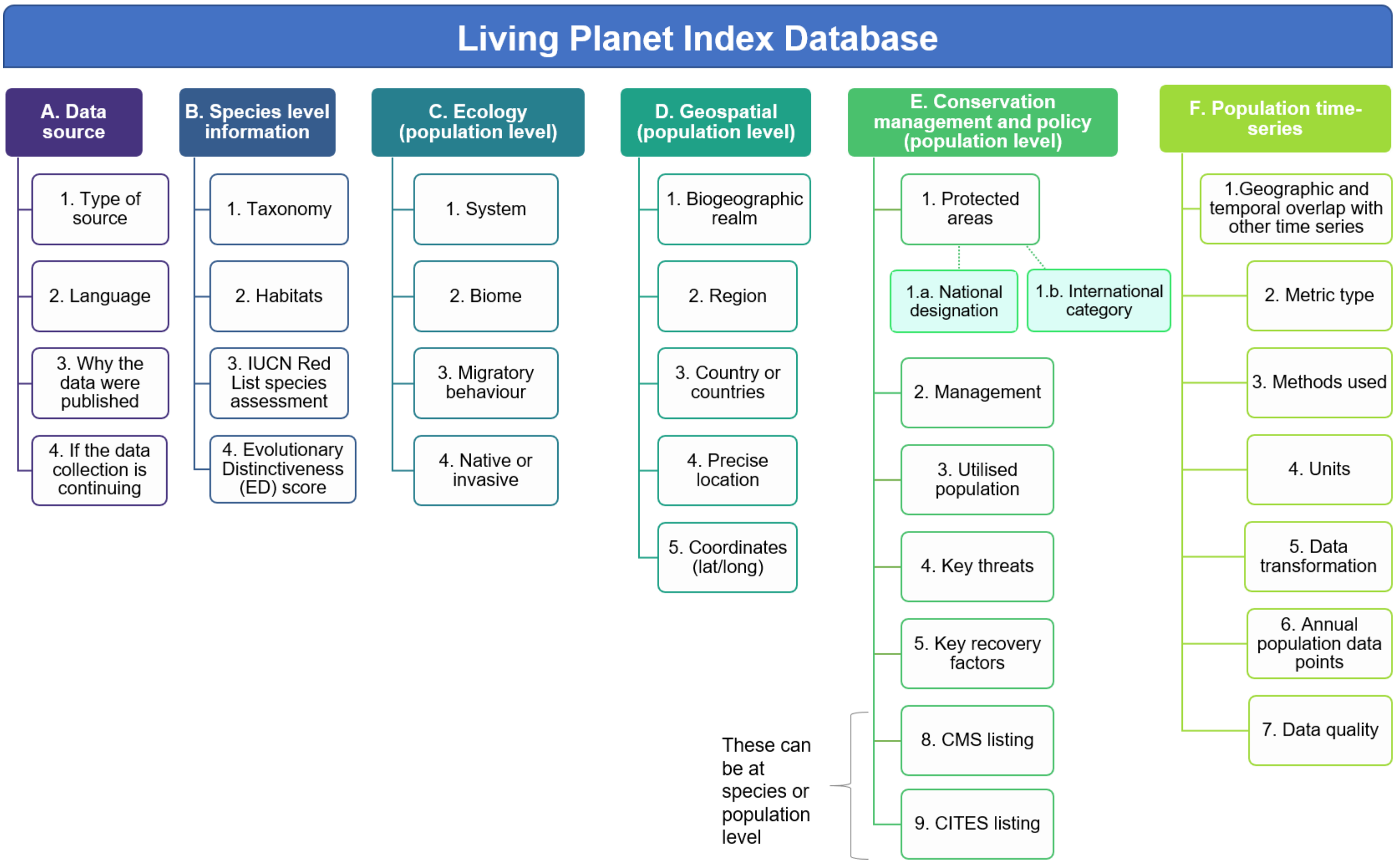


**Supplementary Fig. 5 –** The underlying data of the Living Planet Index database. The data have been grouped thematically (A-F) and the numbered subcategories cover the types of metadata entered for every population within the LPI database. For more information on the definitions and categories see the LPI website supporting documents section (https://www.livingplanetindex.org/supporting_documents).

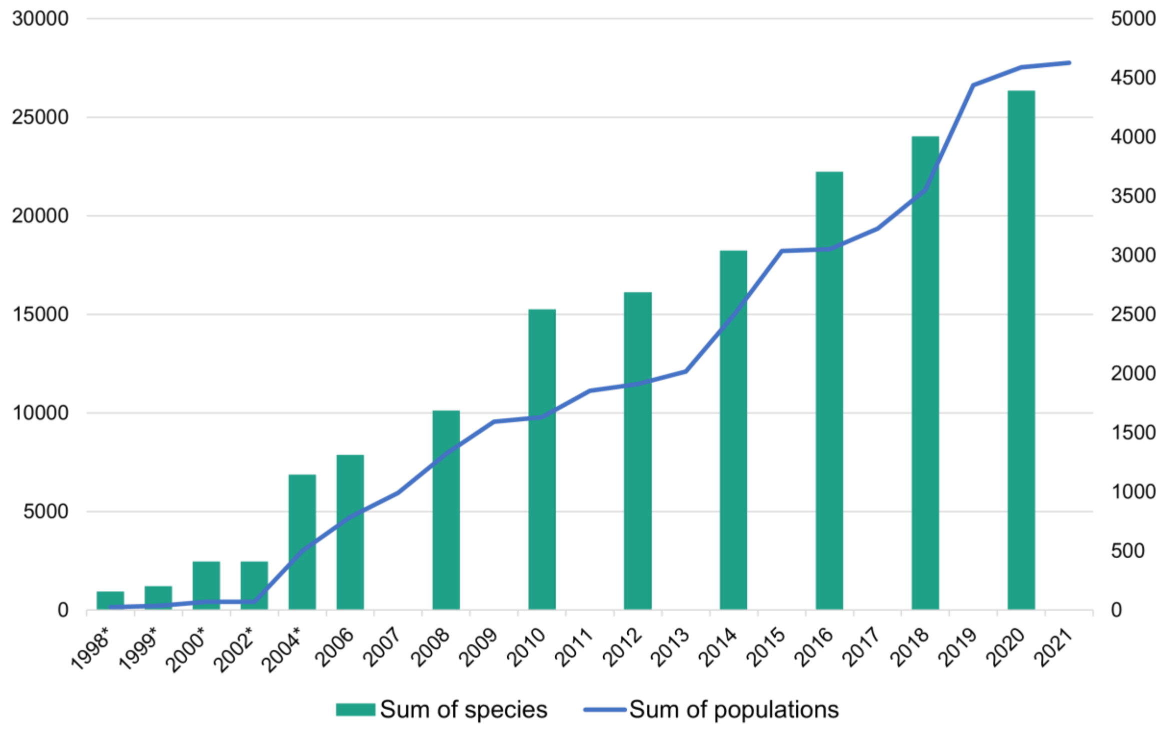


**Supplementary Fig. 6 Growth in populations and species in the Living Planet Database.** Line shows incremental increase in number of populations in the LPD. Bars show number of species in each Living Planet Report. Minimum estimates for numbers of populations were made for years 1998 to 2004 (*)

#### D. Summary of LPD data diagnostics for underrepresented taxa and realms

| Marine priorities ranked by proportional difference | | |
|---|---|---|
| **Rank** | **Realm** | **Taxa** |
| 1 | Tropical and Sub-tropical Indo-Pacific | Fishes |
| 2 | Temperate and Antarctic | Fishes |
| 3 | Atlantic Tropical and Sub-tropical | Fishes |

**Supplementary Table 7. Top 3 marine groups with the highest proportional difference between observed and expected number of species for the LPI dataset in August 2020**. Where observed is the number of species in the dataset and expected is the current estimated number of extant species for that group (see [6] for data sources used).

| Terrestrial priorities ranked by proportional difference | | |
|---|---|---|
| **Rank** | **Realm** | **Taxa** |
| 1 | Neotropical | Fishes |
| 2 | Indo-Malaya | Fishes |
| 3 | Afrotropical | Fishes |
| 4 | Neotropical | Reptiles |
| 5 | Afrotropical | Reptiles |
| 6 | Indo-Malaya | Reptiles |
| 7 | Neotropical | Amphibians |
| 8 | Afrotropical | Amphibians |
| 9 | Indo-Malaya | Amphibians |
| 10 | Palearctic | Fishes |

**Supplementary Table 8. Top 10 terrestrial and freshwater groups with the highest proportional difference between observed and expected number of species for the LPI dataset in August 2020.** Where observed is the number of species in the dataset and expected is the current estimated number of extant species for that group (see [6] for data sources used)

#### E. A visualisation of the data aggregation process for the global LPI method

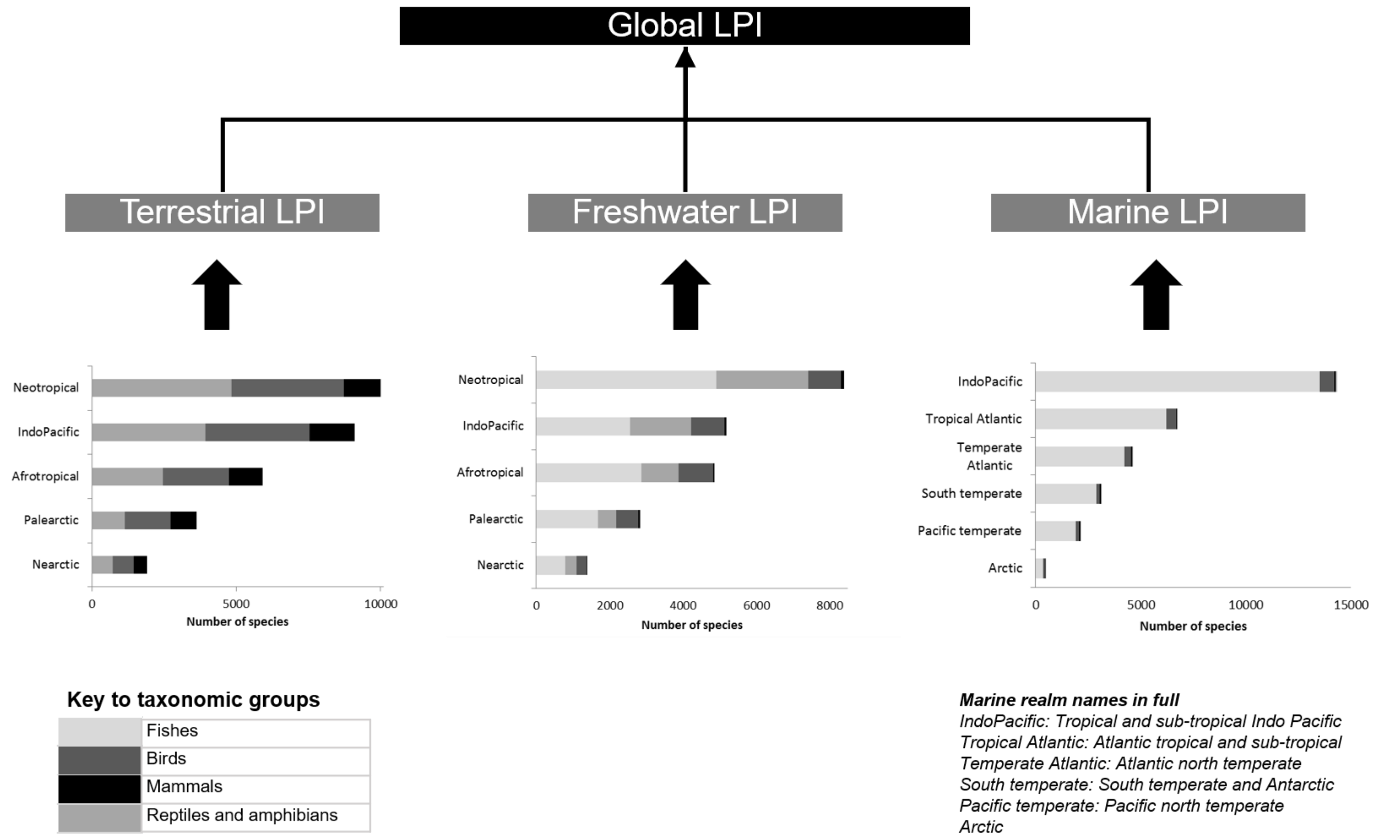


**Supplementary Fig.7. A visualisation of the weighting process for the global LPI**. Figure sourced from: McRae, et al. [6]

2 WWF. Living planet report 2020 - Bending the curve of biodiversity loss. (WWF, Gland, Switzerland, 2020).

3 IPBES. in *Plenary of the Intergovernmental Science-Policy Platform on Biodiversity and Ecosystem Services. Third session.*

4 Collen, B. *et al.* Monitoring change in vertebrate abundance: The Living Planet Index. *Conservation Biology* **23**, 317-327, doi:10.1111/j.1523-1739.2008.01117.x (2009).

5 Loh, J. *et al.* The Living Planet Index: Using species population time series to track trends in biodiversity. *Philosophical Transactions of the Royal Society B: Biological Sciences* **360**, 289-295, doi:10.1098/rstb.2004.1584 (2005).

6 McRae, L., Deinet, S. & Freeman, R. The diversity-weighted Living Planet Index: Controlling for taxonomic bias in a global biodiversity indicator. *PLOS ONE* **12**, e0169156, doi:10.1371/journal.pone.0169156 (2017).

7 Butchart, S. H. *et al.* Global biodiversity: Indicators of recent declines. *Science* **328**, 1164-1168, doi:10.1126/science.1187512 (2010).

8 Tittensor, D. P. *et al.* A mid-term analysis of progress toward international biodiversity targets. *Science* **346**, 241-244, doi:10.1126/science.1257484 (2014).
